# Heterocyclic aromatic amines (HAAs) target mitochondrial physiology

**DOI:** 10.1101/2022.03.17.484822

**Authors:** Shreesh Raj Sammi, Tauqeerunnisa Syeda, Rachel Foguth, Jason Cannon

## Abstract

Heterocyclic aromatic amines (HAA) may be found naturally in plants or are formed through the Maillard reaction when meat is cooked at high temperatures. Previous studies have indicated HAAs are especially toxic to dopaminergic neurons, whereas specific exposures may also affect other neuronal populations such as cholinergic neurons. Biochemical mechanisms of neurotoxic action implicate elevated oxidative stress and afflicted mitochondria. Notably, these mechanisms are of importance in Alzheimer’s disease (AD) and Parkinson’s disease (PD). Most neurodegenerative disease cases are sporadic, where environmental and dietary factors may modulate risk. To further investigate HAA neurotoxicity, we tested the effect of common HAA (harmane, harmine, norharmane, PhIP, and HONH-PhIP) exposure on mitochondrial physiology, a key pathogenic target in AD and PD. Upon assessing mitochondrial bioenergetics, we observed a significant reduction in basal respiration, ATP production, maximal respiratory capacity, and non-mitochondrial respiration, indicating that HAA negatively impacts mitochondrial respiration. Followed by more specific studies on individual mitochondrial complexes, it was found that harmane, harmine, norharmane specifically inhibit complex I enzyme activity, whereas HONH-PhiP, a reactive metabolite of PhIP, inhibits Complex III enzyme activity. Our findings provide significant advancement with respect to underlying mechanisms of toxicity, though also validating HAA exposure as a potential risk factor for AD and PD.

## 1. INTRODUCTION

Alzheimer’s disease (AD) and Parkinson’s disease (PD) are the two most common forms of neurodegenerative disorders affecting nearly 5.8 million (1) and 1 million (2) Americans respectively. Almost 90% of the AD (3) and PD (4) cases are sporadic in nature (with no known cause). While environmental exposures such as pesticides have been extensively studied for etiological roles in these diseases, no single exposure has been found to account for a large number of cases. In contrast to environmental exposures, dietary toxicants remain relatively understudied. The study of dietary neurotoxicants is important to a) determine the underlying pathogenic mechanisms that may influence neurodegeneration and b) identify modifiable risk factors that can limit the onset or progression of disease pathology. Previous studies have identified significant neurotoxic effects of dietary toxicants such as heterocyclic aromatic amines (HAA) on dopaminergic and cholinergic neurons, linking their potential association with AD and PD (5-10). HAAs are produced in meat upon high temperature cooking, through Maillard reaction between amino acids, sugar, and creatinine or pyrolysis of aromatic amino acids (11, 12). HAAs have been classified into different subclasses, α-carbolines, aminoimidazoaazarenes (AIAs), and β-carbolines (9). Amongst AIA HAAs 2-Amino-1-methyl-6-phenylimidazo[4,5-*b*] pyridine (PhIP) is the most abundant and extensively studied HAA (13), where levels in chicken may reach 480 ng/g and human exposures of 0.5 - 1 µg/day (though exposures could be as high as 5-50 µg) (14-17). β-carbolines are also found in plants besides being formed in over cooked meat, fish, brewed coffee and tobacco (20). Most metabolism occurs in liver (21). Previous studies from our lab have identified neurotoxicity of HAAs using various *in vitro, in vivo* and cell-free systems. Acute PhIP exposure induced aberrations in nigrostriatal dopamine system in Sprague-Dawley rats (5). In another study PhIP and its metabolite, HONH-PhIP exhibited selective toxicity towards primary dopaminergic neurons (22). Similarly, selective toxicity of multiple other HAAs was observed in primary midbrain neurons (6). The β-carboline, harmane exhibited dopamine neurotoxicity in *Caenorhabditis elegans*, which was found to be independent of dopamine transporter (a mechanism involved in uptake of known PD toxicants) (8). In addition to the effect on dopaminergic neurons, recent studies on PhIP have also identified neuropathological hallmarks relevant to AD such as an increase in expression of amyloid precursor protein (APP), APP metabolizing enzyme BACE1 and amyloid β aggregation (10). Furthermore, the neurotoxicity of HAAs have been found to be exacerbated in presence of neuromelanin, pointing towards a plausibly heightened neurotoxicity in humans (10, 23). Thus, and coalescing literature suggests that HAAs could be a potential risk factor in both AD and PD due to overlapping mechanisms involved. Given that mitochondrial toxicity is a common characteristic in both AD and PD we aimed to test whether mitochondrial physiology was a direct HAA target, where we focused on mitochondria bioenergetics and respiration. This study identifies critical aspects pertaining to mitochondrial respiration, establishing a link with AD and PD pathology.

## 2. MATERIALS AND METHODS

### 2.1. HAAs

Harmane (#103276), Harmine (286044-1G), Norharmane (# N6252-1G), were purchased from Sigma; PhIP (# A617000), was purchased from Toronto Research Chemicals.

### 2.2. Mitochondrial stress test

Primary cortical neurons were used for assessing mitochondrial function. Primary cortical culture were prepared from the brains isolated from embryonic day 17 pregnant Sprague Dawley rat embryos (Harlan, Indianapolis, IN) using a previously described method (30). Briefly, E17 rat embryos were sacrificed, and the brains were dissected to isolate cortical layers. The tissue was then incubated in papain solution for one hour at 37 degrees. Following digestion, the tissue was mechanically dissociated by vigorous pipetting and resuspended in plating media (Neurobasal supplemented with B27 and 5 % FBS) (30). Cells were plated at a density of 75,000 cells per well in Agilent XFp miniplate. The next day, plating media was replaced with Neurobasal containing B27 alone. Seven days after the plating, the cultures were treated with PhIP or HONH-PhIP at concentrations of 1-10 µM for 24 h. All animal procedures were approved by the Purdue Animal Care and Use Committee.

To assess the effect of HAAs on mitochondrial bioenergetics, a mitochondrial stress test was performed 24 hours after exposure to HAAs using a Seahorse XFp Analyzer and the Seahorse XFp Cell MitoStress test reagents. The mitochondrial stress test determines various aspects of the electron transport chain by measuring the oxygen consumption rate (OCR) and extracellular acidification rate (ECAR) of cells, including changes in basal respiration, ATP production, coupling efficiency, proton leak, maximal respiration, non-mitochondrial respiration, and spare capacity. Oligomycin, FCCP, and Rotenone and Antimycin A are sequentially injected to measure different parameters of mitochondrial function; Rotenone is a potent inhibitor of complex I, Antimycin A is an inhibitor of complex III, Oligomycin is an inhibitor of complex V of the mitochondrial respiratory chain, and FCCP is uncoupler of mitochondrial oxidative phosphorylation (31, 32).

Mitochondrial stress test was performed using Seahorse XFp Extracellular Flux analyzer (Agilent, Santa Clara, CA), according to manufacturer protocol. In brief, one hour before mitochondrial stress test, media was changed to basic media (Agilent base media containing 15.5 mM glucose, 3.76 mM L-glutamine, 1mM sodium pyruvate). Cell plates were then incubated at 37 °C in a non-CO_2_ incubator. Sensor cartridges were hydrated overnight in a non-CO_2_ incubator. Sensor cartridge was loaded with oligomycin, carbonyl cyanide-p-trifluoromethoxyphenylhydrazone (FCCP), and rotenone + antimycin A in the respective ports and injected sequentially to obtain concentrations of 1 μM, 0.5 μM, and 0.5 μM, respectively. Each plate was normalized to control, and each experiment was run 3-4 times. The data generated from Seahorse XFp Extracellular Flux analyzer was plugged in Agilent Seahorse XF Cell Mito Stress Test report generator, and the values were then plotted and analyzed by GraphPad Prism.

HAA doses were 0-10 µM for PhIP and HONH-PhIP and 0-500 µM harmane, harmine and nor harmane. The dose rationale is based upon chosing HAA doses inclusive of the positive control for the assay and an established PD model toxicant and risk factor, rotenone, which is commonly used at 0.5 µM in this specific assay (23, 33, 34).

### 2.3. Specific mitochondrial complex activities

All animal studies were approved by the Purdue Institutional Animal Care and Use Committee (IACUC). Mitochondria were isolated from liver of male Sprague Dawley rats according to the protocol. Spinazzi et al., 2012 (35). Briefly, rats were decapitated, and the liver was isolated and rinsed in liver homogenization medium (LHM) (0.2M mannitol, 50mM sucrose, 10mM KCl, 1mM EDTA, 10mM HEPES, pH 7.4), chopped, and homogenized with a Potter-Elvejem homogenizer. The liver homogenate was centrifuged for 10 minutes at 1,000xg, 4°C, the supernatant was transferred to a new tube and centrifuged for 10 minutes at 3,000xg, 4°C. The supernatant was aspirated, the pellet was resuspended and centrifuged three more times for 10 minutes at 3,000xg, 4°C; the mitochondrial pellet was suspended in phosphate buffer.

Individual mitochondrial complex activities (I, II, III and IV) were measured in the same way as described by Spinazzi et al., 2012 (35). HAA dosing strategy was based upon inclusion of doses relevant to known positive controls in each assay rather than environmental relevance.

Complex I was measured by incubation of 10 μg mitochondria, 3 μg bovine serum albumen (BSA), 0.3 mM KCN, 0.1 mM NADH, 50 μM potassium phosphate buffer (pH 7.5), and 10 μM rotenone, 0-500 μM harmane, 0-500 μM harmine, 0-500 μM nor harmane, 0-10 μM PhIP or 0-10 μM HONH-PhIP for 10 - 15 minutes. Baseline absorbance was measured at 340 nm for 3 minutes prior to the addition of 60 μM ubiquinone; post ubiquinone addition absorbance was measured for another 3 minutes.

Complex II activity was measured by incubation of 1 μg BSA, 0.3 mM KCN, 20 mM succinate, 0.002175% 2,6-Dichlorophenolindophenol (DCPIP), 10 μg mitochondria, 25 mM potassium phosphate (pH 7.4), and 10 mM malonate, 0-500 μM harmane, 0-500 μM harmine, 0-500 μM nor harmane, 0-10 μM PhIP or 0-10 μM HONH-PhIP for 10 - 15 minutes. Baseline absorbance was measured at 600 nm for 3 minutes prior to the addition of 50 μM decylubiquinone; post decylubiquinone addition absorbance was measured for another 3 minutes.

Complex III activity was measured by incubation of 75 μM oxidized cytochrome c, 0.5mM KCN, 0.1mM EDTA, 0.025% Tween 20, 3 μg mitochondria, 25mM potassium phosphate (pH 7.5),and 0.1 μg antimycin A, 0-500 μM harmane, 0-500 μM harmine, 0-500 μM nor harmane, 0-10 μM PhIP or 0-10 μM HONH-PhIP for 10-15 minutes. Baseline absorbance was measured at 550 nm for 3 minute prior to the addition of 0.1 mM decylubiquinol; post decylubiquinol addition absorbance was measured for another 3 minutes Complex IV activity was measured by incubation of 60 μM reduced cytochrome c, 50 mM potassium phosphate (pH 7.0), and 0.3 mM KCN, 0-500 μM harmane, 0-500 μM harmine, 0-500 μM nor harmane, 0-10 μM PhIP or 0-10 μM HONH-PhIP for 10-15 minutes. Baseline absorbance was measured at 550 nm for 3 minutes prior to the addition of 1 μg mitochondria; post mitochondria addition absorbance was measured for another 3 minutes.

Enzyme activity for all complexes were quantified by the following calculation:

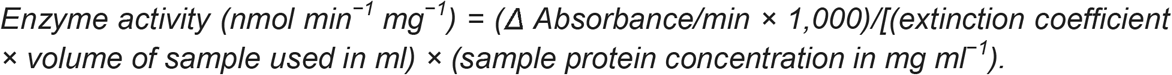

Enzyme activity was expressed as relative enzyme activity, normalized with respect to control. IC50 values were calculated, and curves were plotted using AAT Bioquest for HAAs that inhibited significant (36)

### 2.4. Statistical analysis

Data are presented as mean ± standard error of the mean (SEM). Data was analyzed by one-way ANOVA followed by Dunnet’s post-hoc test for comparison with control in Graphpad Prism 8. p<0.05 was considered statistically significant.

## 3. Results

### 3.1. HAAs exposure alters mitochondrial bioenergetics

Changes in OCR and ECAR levels upon the addition of Oligomycin, FCCP, and Rotenone/Antimycin A in primary cortical neurons treated with PhIP (0-10 µM) or HONH-PhIP (0-10 µM) were assessed. HONH-PhIP induced mitochondrial bioenergetics deficits; PhIP did not alter mitochondrial function, further suggesting the *N*-hydroxylated metabolite is the primary neurotoxicant (***Figure 1***). Alterations in specific bioenergetic parameters were detected following exposure (***Figure 2***). HONH-PhIP (5 and 10 µM) significantly decreased basal respiration (control vs. 5 μM HONH-PhIP : 1.00 ± 0.1975 vs. 0.3110 ± 0.05004, p = 0.0355; control vs. 10 μM HONH-PhIP : 1.00 ± 0.1975 vs. 0.3441 ± 0.06646, p = 0.0308) (***Figure 2***).

**Figure 1:**
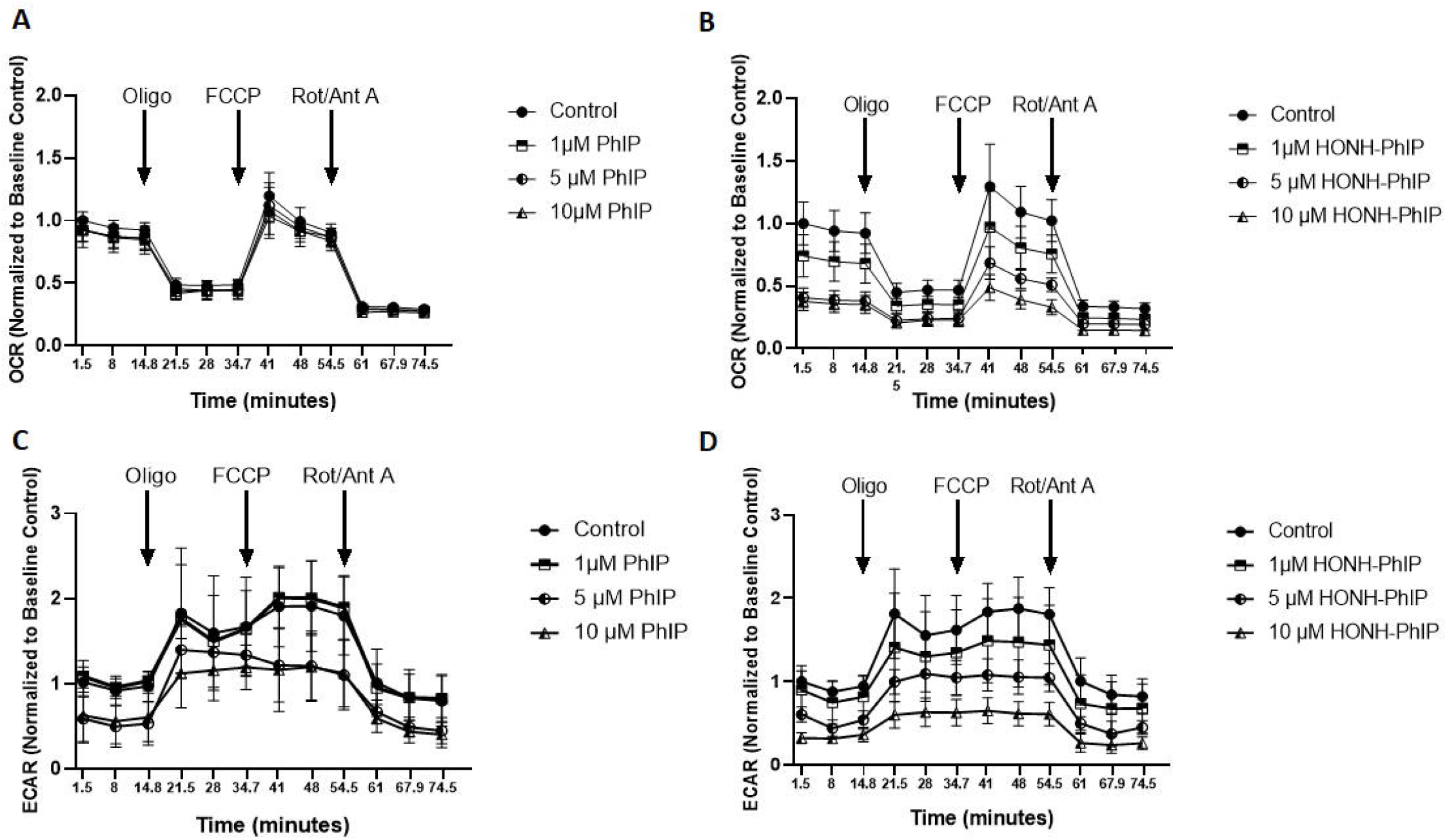
HONH-PhIP significantly decrease mitochondrial respiration after 24 hours of exposure. Primary cortical neurons were treated with A, C) PhIP or B, D) HONH-PhIP for 24 hours. A mitochondrial stress test was then performed on cells and the change in the Oxygen consumption rate (OCR) and extracellular acidification rate (ECAR) was determined using a Seahorse XFp analyzer post sequential addition of oligomycin, FCCP, and Rotenone +Antimycin A, as indicated by arrows. Data are expressed as mean ± S.E.M. Results are representative of 3-4 independent experiments.

**Figure 2:**
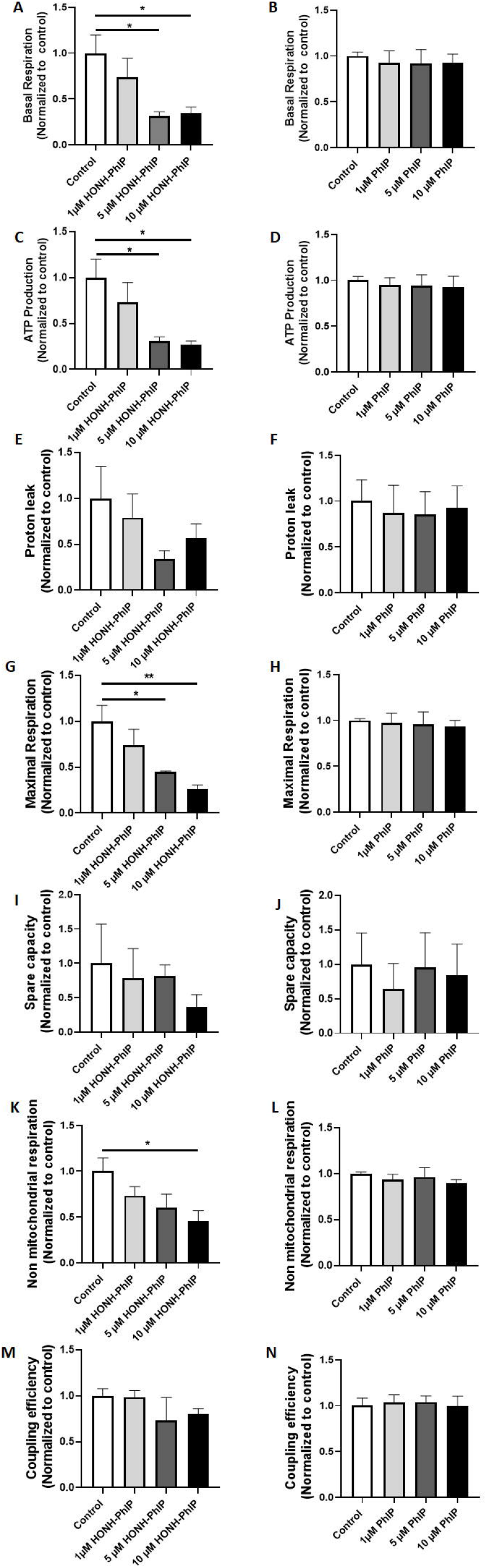
HONH-PhIP significantly decreased Basal respiration, ATP production, maximal capacity and non-mitochondrial respiration. Mitochondrial bioenergetics of primary cortical neurons treated with PhIP (A, C, E, G, I, K and M) or HONH-PhIP (B, D, F, H, J, L, and N) for 24 hours was tested through mitochondrial stress tests. Post treatment, the change in the Oxygen consumption rate levels were determined using a Seahorse XFp analyzer. Basal respiration (A, and B), ATP production (C, and D), proton leak (E and F), maximal capacity (G and H), spare capacity (I and J), non-mitochondrial respiration (K, and L) and coupling efficiency (M, and N) were calculated from the mitochondrial stress tests to determine aspects of decreased respiration. Data are expressed as mean ± S.E.M. Results are representative of 3-4 independent experiments. *p<0.05, **p<0.01.

ATP production was determined by treating with Oligomycin. Oligomycin decreases electron flow through the electron transport chain, resulting in a decrease in OCR, which is linked to ATP production (37). ATP production was significantly decreased on exposure to 5 μM and 10 μM HONH-PhIP (5 and (control vs. 5 μM HONH-PhIP: 1.000 ± 0.2009 vs. 0.3081 ± 0.04577, p = 0.0369; control vs. 10 μM HONH-PhIP: 1.000 ± 0.2009 vs. 0.2715 ± 0.03788, p = 0.0185) (Figure 2).

Maximal respiration capacity was determined by treating with FCCP; electron flow through the electron transport chain is uninhibited, and OCR reaches the maximum. 0.5 μM and 10 μM HONH-PhIP significantly decreased maximal respiration capacity (control vs. 5 μM HONH-PhIP: 1.000 ± 0.1737 vs. 0.4494 ± 0.009668, p = 0.0447; control vs. 10 μM HONH-PhIP: 1.000 ± 0.1737 vs. 0.2655 ± 0.04185, p = 0.0054) (Figure 2).

Non-mitochondrial respiration driven by processes outside the mitochondria. Injection of a mixture of rotenone and antimycin A, inhibits mitochondrial respiration and enables the calculation of non-mitochondrial respiration. HONH-PhIP (10 μM) significantly decreased non-mitochondrial respiration (Control vs. 10 μM HONH-PhIP: 1.000 ± 0.1454 vs. 0.4536 ± 0.1157, p = 0.0402) (***Figure 2***).

No significant change was reported in proton leak, spare capacity, and coupling efficiency on HONH-PhIP exposure (Figure 2).

Data from the MitoStress assay suggest that HONH-PhIP decreases mitochondrial bioenergetics.

### 3.2. Acute HAAs exposure inhibits Specific mitochondrial complex activities in isolated mitochondria

To determine the mechanism of toxicity leading to altered mitochondrial bioenergetics, individual mitochondrial complex activities were measured in mitochondria isolated from rat liver. Mitochondria were acutely treated with harmane, harmine, nor harmane, PhIP, or HONH-PhIP. Harmane, harmine, and nor harmane treatment significantly decreased the activity of complex I in a concentration dependent manner.

Mitochondria treated with 25 µM harmane (0.630 ± 0.118, *p* = 0.0284), 50 µM harmane (0.558 ± 0.036, *p* = 0.0087), 100 µM harmane (0.432 ± 0.077, *p* = 0.0011), 250 µM harmane (0.451 ± 0.065, *p* = 0.0015), and 500 µM harmane (0.194 ± 0.078, *p* < 0.0001) exhibited inhibition of complex I activity in comparison to control (1.000 ± 0.000) (Figure 6A).

Mitochondria treated with 50 µM harmine (0.686 ± 0.091, *p* = 0.0495), 100 µM harmine (0.597 ± 0.071, *p* = 0.0104), 250 µM harmine (0.557 ± 0.049, *p* = 0.0051), and 500 µM harmine (0.437 ± 0.129, *p* = 0.0007) exhibited inhibition of complex I activity in comparison to control (1.000 ± 0.000) (Figure 3B).

**Figure 3:**
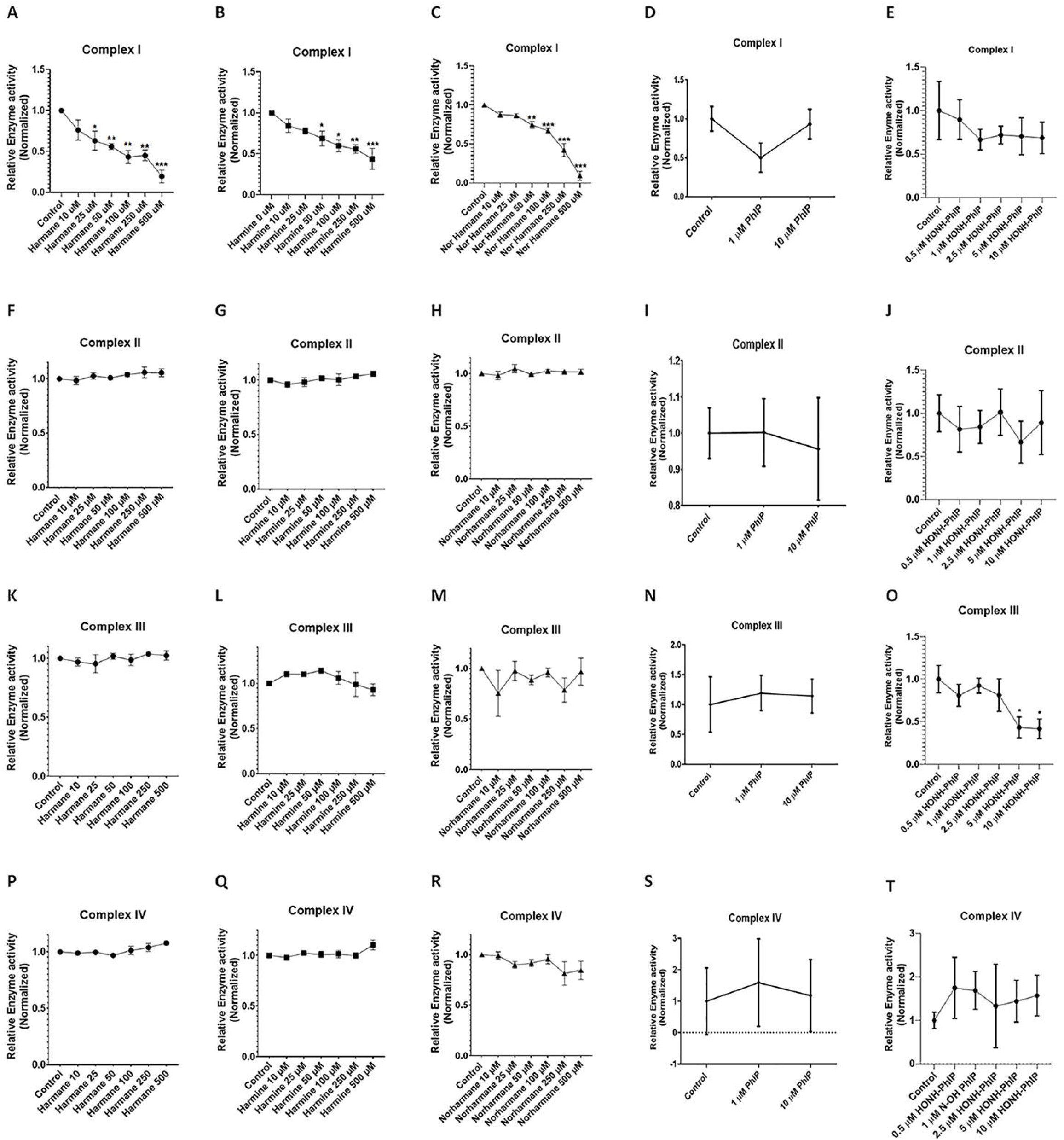
HAAs alter mitochondrial complex activity: Effect of HAAs on mitochondrial complex activity was evaluated on mitochondria isolated from rat liver. Complex I: β-carbolines, Harmane, harmine, Nor harmane exhibited dose dependent inhibition of mitochondrial complex I (A, B, C respectively), while PhIP and NOH-PhIP were devoid of any effect on Complex I (D, E). Complex II: HAAs, Harmane, harmine, Nor harmane, PhIP, and NOH-PhIP did not exhibit any effect on complex II activity (F, G, H, I, J). Complex III: Harmane, harmine, Nor harmane, and PhIP were devoid of any effect on mitochondrial complex III (K, L, M, N respectively), whereas PhIP metabolite NOH-PhIP exhibited dose dependent inhibition of Complex III activity (O). Complex IV: HAAs, Harmane, harmine, Nor harmane, PhIP, and NOH-PhIP did not exhibit any effect on complex IV activity (P, Q, R, S, T). Data analyzed using one-way ANOVA followed by Dunnett’s post hoc test. *p<.05, **p<.005, and ***p<.001 (n =3).

Mitochondria treated with, 50 µM nor harmane (0.744 ± 0.043, *p* = 0.0069), 100 µM nor harmane (0.668 ± 0.032, *p* = 0.0008), 250 µM nor harmane (0.420 ± 0.082, *p* < 0.0001), and 500 µM nor harmane (0.091 ± 0.057, *p* < 0.0001) exhibited inhibition of complex I activity in comparison to control (1.000 ± 0.000) (Figure 3 C).

PhIP and NOH-PhIP lacked any effect on complex I activity (***Figure 3 D, E***). The above results showed affliction of complex I activity in response to β-carbolines.

Complex II activity remained unaltered upon treatment with any HAAs (***Figure 3 F, G, H, I, J***).

Similarly, complex III activity remained mostly unaltered (***Figure 3 K, L, M, N***) except for mitochondria treated with HONH-PhIP which exhibited a significant reduction in Complex III activity at doses 5 µM HONH-PhIP (0.4320 ± 0.1222, *p* = 0.0245), and 10 µM HONH-PhIP (0.4164 ± 0.1140, *p* = 0.0203) in comparison to control (1.000 ± 0.1598) (***Figure 3O***).

Treatment with HAAs did not exhibit any effect on mitochondria complex IV (***Figure 3 P, Q, R, S, T***). The above results showed that HAAs subclass operate with the distinct mechanism. While β-carboline afflict complex I, effect of HONH-PhIP was mostly limited to complex III. IC50 values were calculated and curves were plotted as shown in ***Table 1*** and ***Figure 4***.

**Table 1:** IC 50 of mitochondrial complex inhibition.

| HAA | Complex I | Complex II | Complex III | Complex IV |
| --- | --- | --- | --- | --- |
| **Harmane** | 76.575 μM | N.D. | N.D. | N.D. |
| **Harmine** | 309.327 μM | N.D. | N.D. | N.D. |
| **Nor harmane** | 148.378 μM | N.D. | N.D. | N.D. |
| **PhIP** | N.D. | N.D. | N.D. | N.D. |
| **HONH-PhIP** | N.D. | N.D. | 5.373 μM | N.D. |
| **6-hydroxydopamine** | 10 μM (52) | N.D. | N.D. | 34 μM (52) |
| **Antimycin** | N.D. | N.D. | 22 μM (53) | N.D. |

N.D. = Not determined

**Figure 4:**
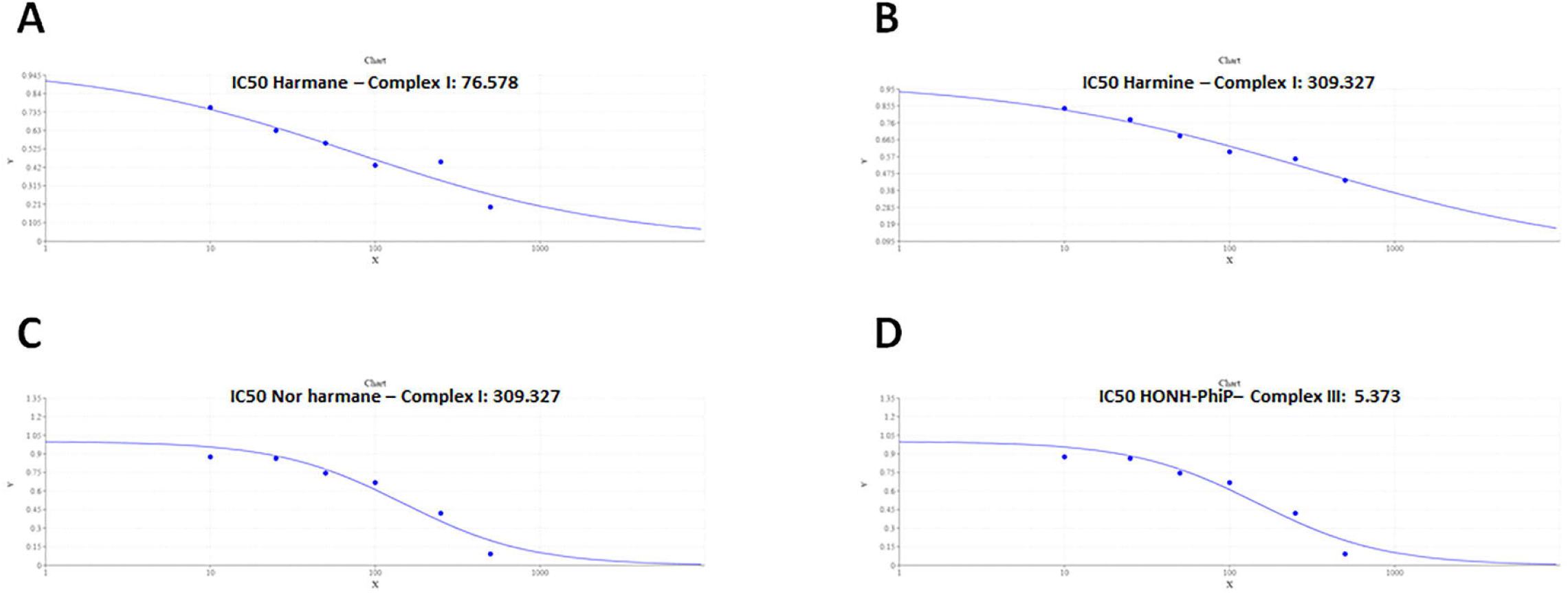
IC50 for Complex I for harmane, harmine, and norharmane and Complex III for HONH-PhIP. Curve and IC50 - Complex I for harmane (A), harmine (B), norharmane (C); Complex III for HONH-PhIP (D).

## 4. Discussion

Mitochondria have been implicated as primary targets in neurodegenerative diseases such as AD and PD. Mitochondrial dysfunction in the form of altered mitochondrial morphology, reduced bioenergetics capacity, compromised enzyme complexes in the Krebs cycle, reduced cytochrome c oxidase activity, and increased oxidative stress is observed in AD and PD. For example, our group has shown that HAA exposure altered mitochondrial bioenergetics, elevated the mitochondrial superoxide levels and mitochondrial fission, reduced healthy mitochondria and mitochondrial membrane potential in SH-SY5Y neuroblastoma cells overexpressing tyrosinase (23). A dose-dependent loss of mitochondria was observed in C. elegans treated with harmane (8). Previous studies showed that harmane derivatives cause mitochondrial dysfunction when injected into the substantia nigra (40, 41). Importantly, relative to neurotoxic exposures, relatively few studies have implicated the specific mitochondrial targets that may underlie resultant bioenergetic deficits. Thus, in the present study, we used both broad markers of cellular bioenergetics and specific mitochondrial respiration assays to link mitochondrial respiration deficits to specific respiration complex inhibition.

Our study showed that mitochondrial function is susceptible to HAAs toxicity, and several aspects of mitochondrial bioenergetics are targeted by HAAs. Using a Seahorse Extracellular Flux Analyzer, we monitored cellular OCR and ECAR as a measures of mitochondrial respiration and glycolysis. OCR reflects mitochondrial function and mitochondrial oxidative phosphorylation (42).

Our data demonstrated that HONH-PhIP (reactive metabolite of PhIP) altered mitochondrial bioenergetics. Treatment with 5 µM and 10 µM HONH-PhIP decreased basal respiration which is controlled by ATP turnover and partly by proton leak and substrate oxidation (37). HONH-PhIP 5 µM and 10 µM decreased ATP production, which was measured from the decrease in respiration on inhibiting the complex V with oligomycin.

Treatment with HONH-PhIP 5 µM and 10 µM dose decreased the maximum respiratory capacity (caused by addition of FCCP). A decrease in maximum respiratory capacity is an indicator of potential mitochondrial dysfunction (37). Primary cortical neurons treated with 10 µM HONH-PhIP had lower non-mitochondrial respiration.

These data indicate that HONH-PhIP decreases mitochondria bioenergetics; In order to further delve into the underlying mechanisms, we tested the effect of HAAs on individual mitochondrial complex activity. We observed a distinct mechanistic pattern amongst HAA subclasses, AIA and β-carbolines. A significant dose dependent inhibition in complex I activity was observed in case of β-carbolines (harmane, harmine and norharmane). Conversely, AIA, HONH-PhIP was devoid of any effect on complex I activity. However, a significant inhibition of complex III activity was observed upon treatment with HONH-PhIP.

While complex I deficits have been extensively studied in PD, it is worth noting that HAA induced inhibition of Complex I and complex III activity are relevant to both AD and PD (43-45). Complex I and complex III are considered to be the primary source of reactive oxygen species in physiological and pathological processes. Lowered complex I activity in platelet mitochondria has been seen in early untreated PD patients (46). PD relevant mitochondrial toxins MPTP (47, 48) and rotenone (34, 49), selectively inhibit complex I and cause PD-like clinical features. Additionally, subunits of complex I (NDUFA4 and NDUFA9) and Complex III core protein were altered in AD patients (50, 51). Previous studies on harmane have exhibited a decrease in functional mitochondria in *C.elegans*, potentially through inhibition of specific mitochondrial complexes, as treatment with mitochondrial complex I activator partially rescued harmane-induced neurodegeneration (8). The current data on complex I inhibition validated the previous findings about harmane led neurodegeneration in *C. elegans* (8).

While the current studies have utilized higher concentration of HAAs, than what has been found in food, it is noteworthy that in the absence of any guidelines, humans are continuously and chronically exposed to these toxicants rendering HAAs to potentially modulate pathology of neurodegenerative disorders over the span of time. Noteworthy, AD and PD are age dependent disorders with etiology ranging to decades.

Collectively, our data is suggestive of a potential role of HAA induced mitochondrial dysfunction in culminating neurotoxicity, implying that HAA led mitochondrial toxicity could contribute to the advent or progression of neurodegenerative diseases.

## 5. Acknowledgments

HONH-PhIP was a kind gift from Dr. Robert Turesky (University of Minnesota). We thank the laboratory of Dr. Jean-Christophe Rochet for assistance with primary rat cortical cultures.

## 6. Funding

This research was funded by the National Institute of Environmental Health Sciences at the National Institutes of Health (ES025750 to J.R.C. and K99ES032488 to S.R.S).

## Notes

### Competing Interest Statement

The authors have declared no competing interest.

